# Early immune response to *Coccidioides* is characterized by robust neutrophil and fibrotic macrophage recruitment and differentiation

**DOI:** 10.1101/2024.08.21.609001

**Authors:** Nadia Miranda, Oscar A. Davalos, Aimy Sebastian, Margarita V. Rangel, Nicole F. Leon, Bria M. Gorman, Deepa K. Murugesh, Nicholas R. Hum, Gabriela G. Loots, Katrina K. Hoyer, Dina R. Weilhammer

## Abstract

Coccidioidomycosis, or Valley fever, is an emerging respiratory disease caused by soil dwelling fungi of the *Coccidioides* genus that is expected to spread from the southwest into the central U.S. by 2050. While 60% of infections are asymptomatic, the other 40% of patients experience a range of symptoms, from self-limiting pneumonia to life-threatening disseminated disease. The immunological events that underlie the progression to severe disease remain under defined. Here, we probed the early immune response to *Coccidioides* using a high dose of an attenuated strain of *C. posadasii* in a mouse model of infection coupled with single-cell RNA sequencing. At 24 hours post-infection, robust immune infiltration is detected in the lung, marked by high levels of inflammatory PD-L1^+^ neutrophils and fungal-contact dependent pro-fibrotic Spp1^+^ macrophages. These findings elucidate the early dynamics of the host response to *Coccidioides* and provide a deeper understanding of host-pathogen interactions in the lung.

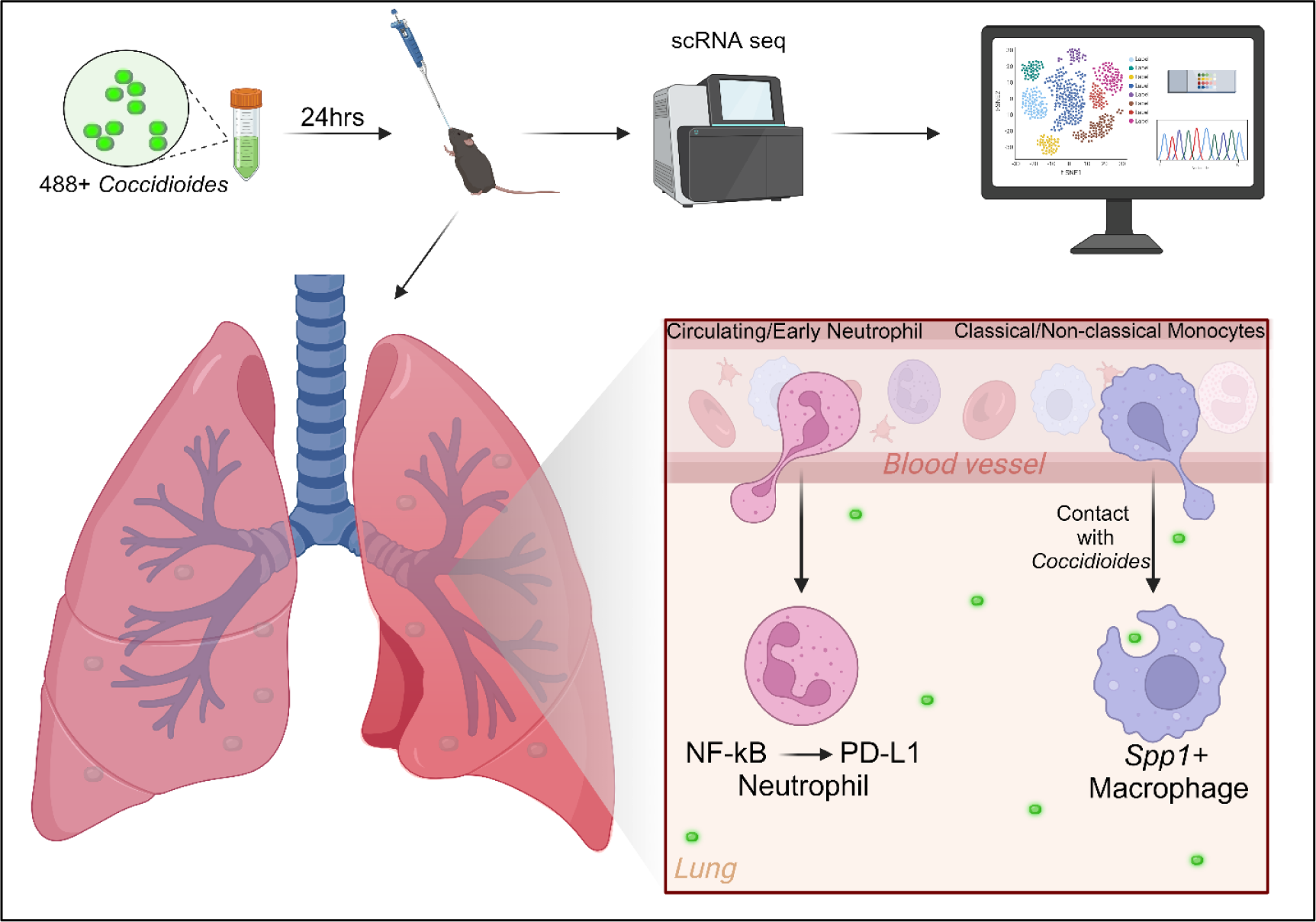

**Importance:** By examining early immune dynamics in the lungs, we uncover critical insights into how myeloid cells, particularly neutrophils and macrophages, are recruited and differentiated during *Coccidioides* infection. The discovery of specific immune cell subsets, such as PD-L1^+^ neutrophils and Spp1^+^ macrophages, which are associated with inflammation and fibrosis, highlights potential targets for therapeutic intervention. These findings provide a deeper understanding of the host-pathogen interactions that occur during *Coccidioides* infection, offering valuable directions for developing more effective treatments and preventive strategies against this increasingly prevalent disease.

## Introduction

Climate change is accelerating the spread of fungal infections due to rising temperatures and shifting environmental conditions (*1*). These changes are expanding the habitats of fungi, transforming previously non-endemic areas in the central United States into new hotspots for infections, thereby increasing the risk to humans, pets, and livestock (*2*). This expanding geographic range of fungi highlights the urgent need for preventative measures to combat the rising threat of fungal diseases. One such emerging fungal infection is *Coccidioides*, an airborne pathogen that is progressively spreading from the southwest towards the central United States due to climate change (*3*).

Coccidioidomycosis, often termed Valley fever, is caused by inhaling the fungi *Coccidioides immitis* or *Coccidioides posadasii* in airborne dust (*3*). *Coccidioides* infections continue to rise each year in California and Arizona, states with mandatory reporting, while disease is underestimated in other southwestern states where reporting is not mandatory. The Center for Disease Control estimates that 500,000 cases are reported every year (*4*). Efforts to reduce the impact of Valley fever have been limited due to the lack of a definitive and rapid diagnostic, an insufficient understanding of the factors that predict which patients will develop severe, even fatal, disease, and a lack of effective therapies for those who experience disseminated disease. Thus, there is an urgent need to define effective host immunity to develop protective vaccines and immune modifying therapeutics, as well as identify diagnostic targets.

The mechanistic details that underlie a protective immune response against *Coccidioides* infection, knowledge needed for effective diagnosis, treatment, and vaccine development, remain under described. The anti-fungal immune response is controlled by the early activation of innate immune cells through pathogen recognition receptors, including Dectin-1/2 and mincle in response to *Coccidioides* antigens, such as spherical outer wall glycoprotein (SOWgp), β-glucans, and chitins (*5, 6*). Next, neutrophils rapidly accumulate within the lung and fungal-induced granulomas, emphasizing their critical role in the early *Coccidioides* and other anti-fungal immune responses (*7–21*). Monocytes recruited from circulation rapidly differentiate into macrophages upon migrating into infected regions of the lung where they produce tumor necrosis factor-alpha (TNF-α), interleukin-6 (IL-6) and reactive oxygen species in response to *Coccidioides*, key hallmarks of an innate inflammatory response (*22, 23*). A coordinated effort between barrier epithelial cells and other innate cells interacting with the pathogen drives the differentiation of effector populations with unique functions specific to the invading pathogen (*24*). While these events are known for some viral, fungal, and bacterial pathogens, the complex and dynamic nature of the immune response to infection with *Coccidioides*, and how the balancing inflammatory functions can result in a managed infection, is not yet known.

This study utilized single-cell RNA-sequencing (scRNAseq) and fluorescently labeled fungal spores to reveal the immediate immune response within the lung elicited against a high dose of an attenuated *Coccidioides* strain. Within 24 hours post-infection (hpi) there is a large influx of neutrophils and macrophages into the lungs. Diverse populations of neutrophils emerge, including a population that expresses high levels of *Cd274,* the gene that encodes PD-L1. Monocyte-derived Spp1^+^ macrophages differentiate in a fungal contact-dependent manner and express a pro-fibrotic gene signature. Labeled *Coccidioides* preferentially associate with neutrophils and Spp1^+^ macrophages, as revealed by tissue imaging and flow cytometry. This multi-omics approach, incorporating flow cytometry, sequencing, and fungal imaging, provides novel insights into the early immune response against *Coccidioides*, offering promising avenues for further exploration in combating airborne infections and developing effective vaccine strategies.

## Results

### Single-cell RNA sequencing reveals robust myeloid cell infiltration into the lungs concurrent with shifts in non-immune cells

To elucidate early events in *Coccidioides* infection, we used scRNAseq to profile the transcriptional response in the lung in both immune and non-immune cells. C57BL/6 mice were intranasally infected with *C. posadasii* Δ*cts2*/Δ*ard1*/Δ*cts3*, and lungs were harvested 24 hpi, along with mock infected controls (**Figure 1A**). Unsupervised clustering of 16,265 and 9,990 cells from uninfected and infected lungs resulted in 12 clusters that were each assigned to a putative cell-type identity based on their unique profile of differentially expressed genes (DEGs) encoding cell-type specific markers (**Figure 1B, D, S1A**). The relative proportion of several immune cell populations increased upon infection, most notably neutrophils and monocytes/macrophages, along with a reduction in the relative proportion of non-immune cells such as endothelial cells, epithelial cells, and fibroblasts (**Figure 1C**). To further understand cellular interactions, CellChat was utilized to examine cellular communication and signaling pathways involved during infection (*25*). Within uninfected lungs, endothelial cells were the dominant cell type in terms of incoming and outgoing signaling strength (**Figure 1E**). Consistent with their increased abundance in the lung, neutrophils and monocyte/macrophages dominate cellular communication in the infected lungs, with a notable shift in signaling strength in both the incoming and outgoing interactions as well as an increase in the number of interactions (**Figure 1E**), suggesting significant reorganization of cellular communication networks in the lung in response to the robust neutrophil and macrophage infiltration.

**Figure 1:**
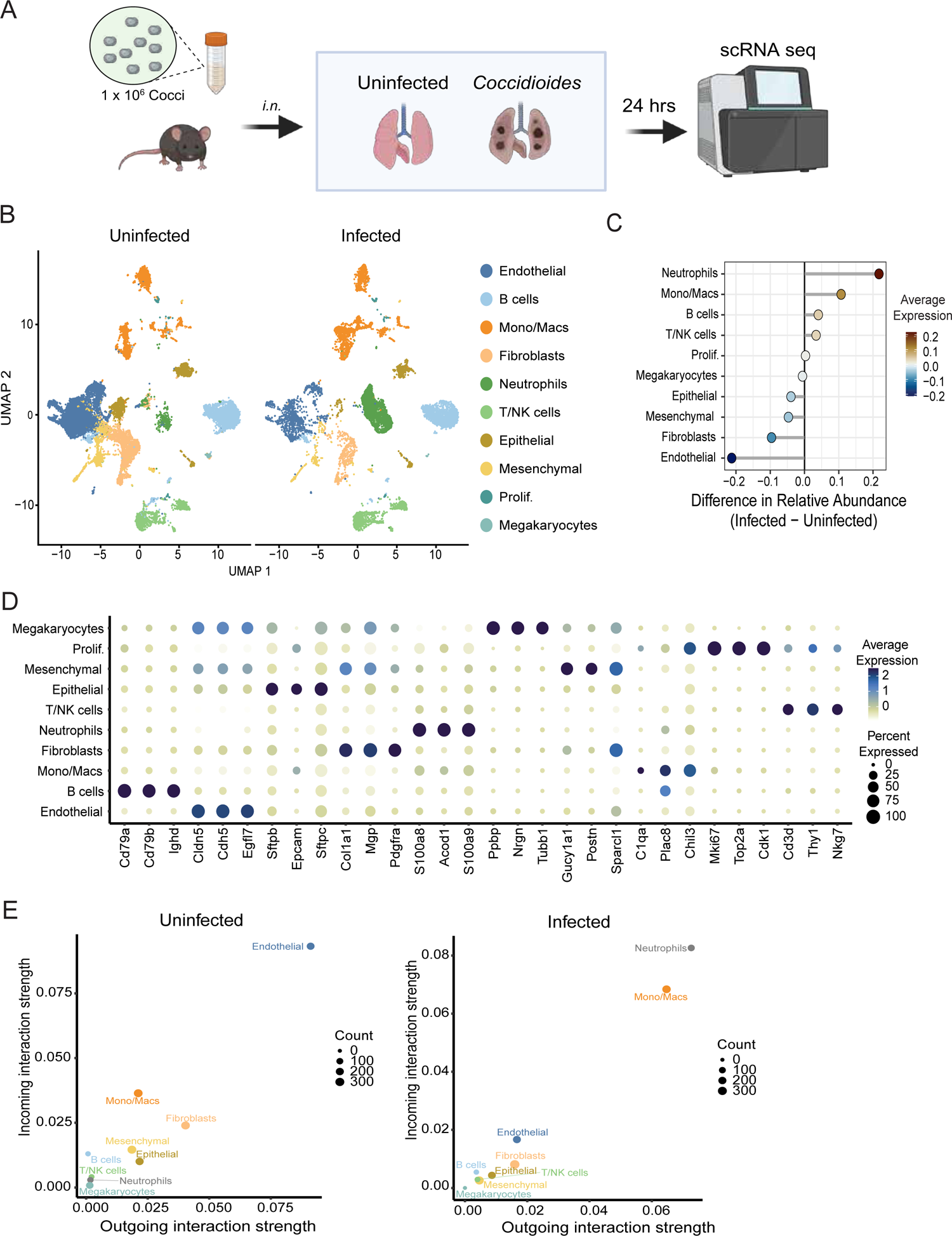
Single cell RNA sequencing reveals shifts in cell populations and immune infiltration in the lungs following C. posadasii infection. (A) Experimental design. Mice were infected intranasally with 1×106 C. posadasii (Δcts2/Δard1/Δcts3) arthroconidia or mock infected. Lungs (n=3 pooled per condition) were harvested for scRNAseq analysis 24 hpi. (B) Uniform Manifold Approximation and Projection (UMAP) visualization of cell clusters identified in each experimental group. (C) Relative cellular proportions in infected versus uninfected lungs. (D) Dot plot showing the expression of select cell-type markers. Dot size represents the fraction of cells of the indicated cluster expressing the markers and the intensity of color represents the average marker expression level in that cluster. (E) CellChat analysis of incoming and outgoing signal strength in the uninfected and infected groups from cell populations in (B). Dot size indicates the number of interactions. n=3 pooled per condition for sequencing

Endothelial, epithelial, fibroblast, and mesenchymal lung cells maintain homeostasis within the lung microenvironment and are among the first cell types to encounter invading pathogens. To probe the response of these resident lung cells to infection, we subclustered the non-immune cells, resulting in 18 distinct cell clusters (**Figure 2A** and **B, S1B, S1C**). Although the relative abundance of all non-immune cell types decreased with infection (**Figure 1C**), epithelial cells increased in abundance relative to other non-immune cell types, while endothelial cells decreased (**Figure 2C**). The increase in epithelial cells was primarily due to an expansion of AT2 and ciliated cells, while club and goblet cells remained the same and AT1 cells decreased. MANC fibroblasts increased slightly in abundance while myofib remained steady and lipofib decreased. AEC were the only endothelial subtype to increase slightly while other subtypes either remained steady or decreased. The proportion of all mesenchymal cell subtypes remained steady. These data are summarized in **Figure 2D**.

**Figure 2:**
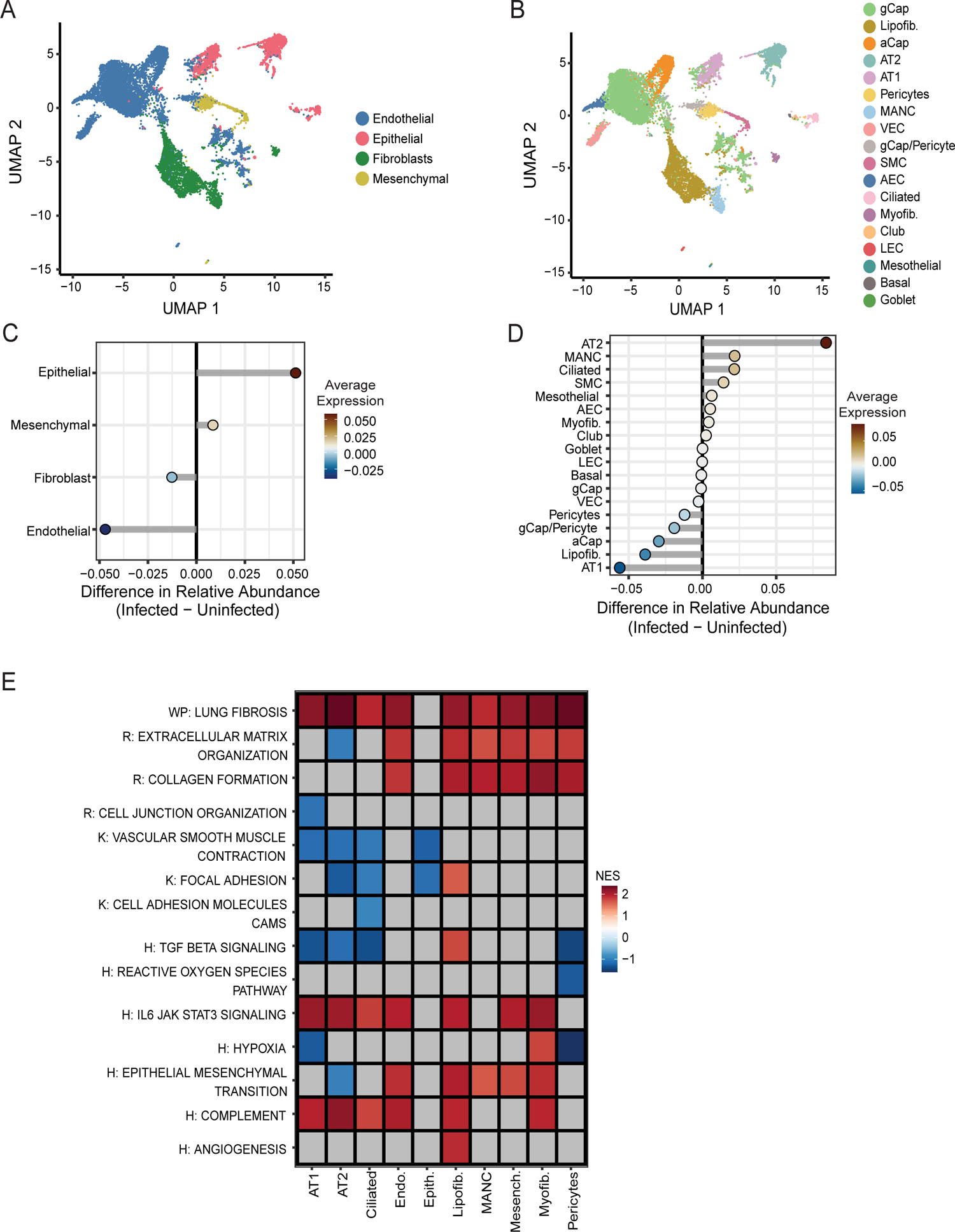
Non-immune cells respond to infection by upregulation of select inflammatory and pro-fibrotic. pathways (A) Integrated UMAP of the main non-immune cell types of the lung. Cells are color-coded by type. (B) Integrated UMAP of subclustered non-immune cell types of the lung. Cells are color-coded by type. Relative cellular proportions of main non-immune cell types (C) and subclustered non-immune cell types (D) in infected versus uninfected lungs. (E) Heatmap of pathway activities scored per cell by AUCell. Enrichment is represented by a scaled average AUCell score per cell type, with blue indicating lower enrichment and red indicating higher enrichment. The X-axis denotes the cell type, color coded by non-immune cell subtype as in (A), and the Y-axis denotes the pathway.

To gain deeper insights into the pathways active within the non-immune subpopulations, we performed an enrichment analysis by identifying differentially expressed genes in infected cells verses uninfected. Strikingly, several pathways involved in fibrosis, including Lung Fibrosis, Collagen Formation, Epithelial Mesenchymal Transition, and Extracellular Matrix Organization pathways were enriched widely across many cell types, including fibroblasts, endothelial cells, mesenchymal, and some subsets of epithelial cells (**Figure 2E**). IL-6/JAK/STAT3 Signaling and Complement were also enriched across multiple cell types. These findings highlight a shift towards profibrotic and inflammatory pathways within non-immune cells in response to infection.

### Diverse neutrophil populations respond to infection

Neutrophils were highly enriched in infected samples (**Figure 1C**), and a significant increase in neutrophils within the lungs of infected mice was additionally confirmed by flow cytometry (**Figure 3A**). To probe the diversity of the neutrophil (neu) response, 2021 cells were extracted and re-clustered, revealing five distinct subtypes: Circulating neu (*Cxcr4*, *Sell*, *Fgl2*), Programmed death-ligand 1 (PD-L1) neu (*Cd274*, *Ccl3*, *Tnf*), NF-кB high neu (*Nfkb1*, *Cxcl3*, *Nr4a3*), Early neu (*Mmp8*, *Retnlg*, *Ngp*), and interferon-stimulated gene (ISG) expressing neu. (*Rsad2*, *Ifit3*, *Ift3b*) (**Figure 3B**, **S2 A-C**). Circulating neu were the predominant subtype in the lungs of uninfected mice, and a dramatic shift towards PD-L1 and NF-кB neu was observed following infection (**Figure 3C**, **S2B**). Gene ontology (GO) analysis demonstrated that neutrophil subpopulations were enriched for distinct pathways (**Figure 3E**, **S2D**). Circulating, Early, and NF-кB neu were enriched for pathways associated with Chemotaxis, Migration, and Cell-Cell Adhesion, with early neu further enriched for pathways associated with Response to Reactive Oxygen Species/Oxidative Stress and NF-кB neu demonstrating enrichment for Cellular Activation/Signaling pathways (**Figure 3E**, **S1D**). ISG neu were distinctly enriched for pathways associated with Response to Viruses and PD-L1 neu demonstrated overlapping enrichment characteristics with NF-кB and ISG neu, enriched primarily in pathways related to Cell Activation and Signaling (**Figure 3E**, **S1D**). Consistent with a shift towards a more activated phenotype upon infection, an increase in TNF and Complement signaling between neutrophils and several immune and non-immune cell types was observed, most notably with monocyte/macrophages (**Figure S2E and S3A**).

**Figure 3:**
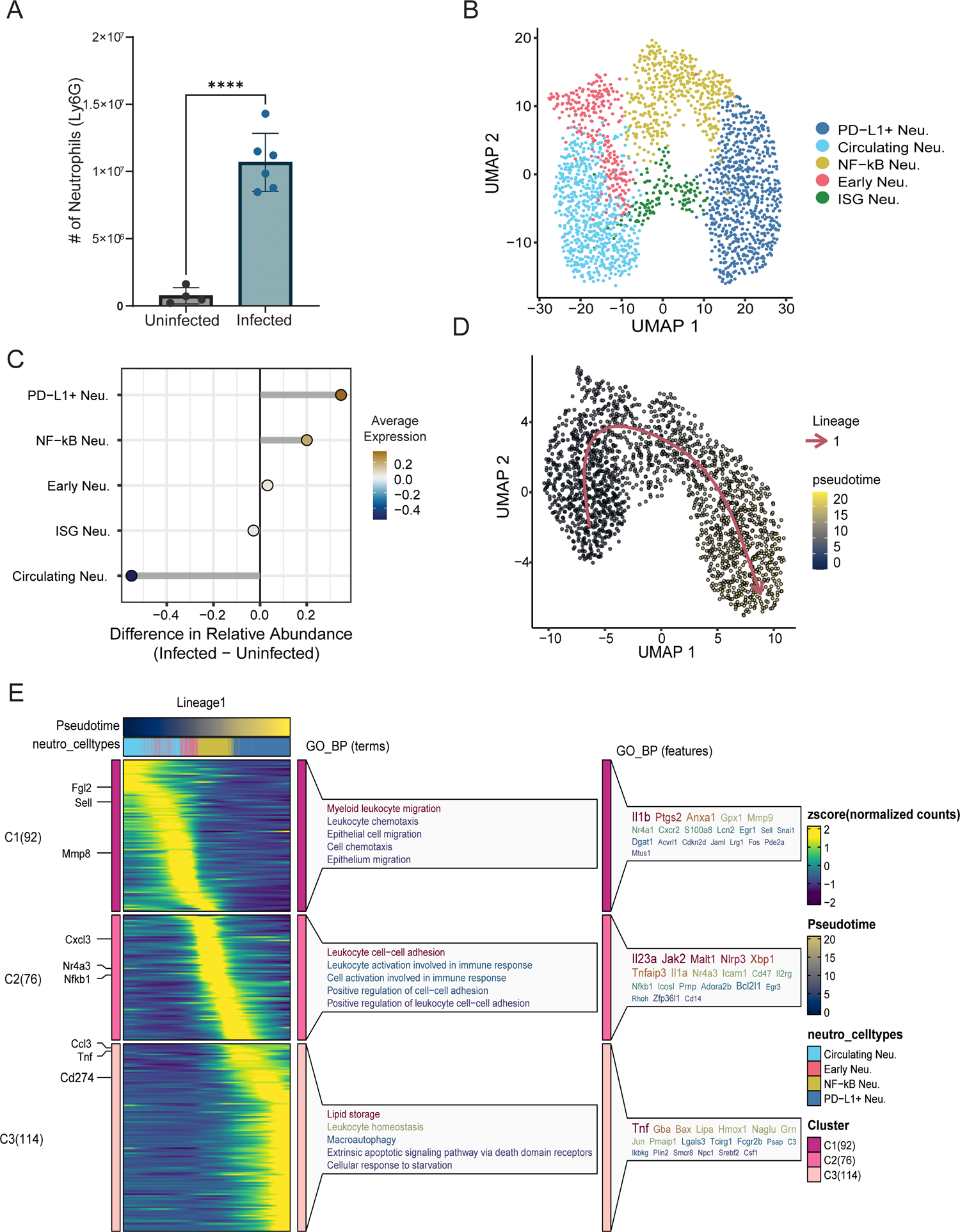
Diverse neutrophil populations accumulate in the lung 24 hpi. (A) Number of neutrophils in the lungs as identified by flow cytometric analysis. (B) Integrated UMAP of subclustered neutrophils. Cells are color-coded by type. (C) Relative cellular proportions of neutrophil subsets. (D) UMAP plot displaying pseudotime trajectory. Cells are color-coded based on their pseudotime values, ranging from blue (early) to yellow (late). The pseudotime trajectory was inferred in an unsupervised manner using Slingshot. (E) Heatmap illustrating the scaled fitted gene expression values along pseudotime. Expression levels are color-coded from low (purple) to high (yellow). The top row for lineage represents pseudotime, ordered from left to right and color-coded based on pseudotime values, ranging from blue (early) to red (late). The next row contains cell type colors, where each bar represents a cell belonging to a specific cell type along the pseudotime trajectory. Rows represent genes, with selected genes highlighted on the left-hand side. Genes were clustered based on fitted expression values, with each cluster represented by row-wise breaks in the heatmap. Each cluster is labeled on the left-hand side by “C#” followed by the number of genes in that cluster in parentheses. Different gene clusters can be distinguished by the clusters on the left-hand side of the heatmap, represented by a color block. Genes from each cluster were then analyzed for gene ontology (GO) enrichment, with the top **5 enriched terms for each cluster shown on the right side of the heatmap. ****p<.00001**

To better understand the differentiation of infection-induced neutrophil subsets along with genes changed over the lineage, we leveraged Slingshot (*26*) to perform a pseudotime trajectory analysis. Initial analysis identified two neutrophil lineages (data not shown). However, as ISG neutrophils were largely unchanged during infection (**Figure 3C**), this population was excluded from subsequent analysis. One neutrophil differentiation lineage was identified, indicating that neutrophils differentiated in a trajectory through Circulating, Early, NF-кB-expressing, and ultimately into PD-L1^+^ neutrophils (**Figure 3D**). Gene expression patterns clustered into three groups and GO analysis of these clusters demonstrated different biological activities associated with each group (**Figure 3E S2D)**. Clusters 1 and 2 (C1 and C2) contained Circulating and Early neu, showing enrichment for migration and IL-1β responses. C2 also demonstrated enrichment for cell-cell adhesion, activation, and IL-23 and was composed of primarily of NF-кB neu. C3 displayed high expression of TNF, PD-L1, Lipid Storage, and Macroautophagy, marking the termination of the neutrophil lineage at PD-L1^+^ neutrophils. (**Figure 3D** and **E**). These data indicate a dynamic neutrophil response occurring early following high dose *Coccidioides* infection.

### Monocyte derived Spp1^+^ macrophages exhibit fibrotic state

Monocyte/macrophage cells were also highly enriched in infected samples. To gain deeper insights into their expression profiles, the Mono/Mac cluster (**Figure 1B**), totaling 2599 cells, was extracted and re-clustered, resulting in five subclusters: Alveolar macrophages (AM; *Plet1, Krt19, Lpl*), Classical monocytes (C. Mono; *Plac8, Ly6c2, Sell*), Non-classical monocytes (NC. Mono; *Adgre4, Ace, Treml4*), Spp1^+^ macrophages (Spp1^+^ Mac; *Fn1, Spp1, Inhba*), MHC II high interstitial macrophages (MHCII IM; *Cd163, H2-Ab1, C1qa*), and Interferon high monocytes (IFN Mono; *Rsad2, Ifit2, Ifit3*) (**Figure 4A** and **Supplemental Figure 3D**). The Spp1^+^ macrophages represent a novel subclass that emerge only upon infection and display the highest relative abundance within infected lungs (**Figure 4B**). In response to *Coccidioides* infection, Spp1^+^ macrophages express high levels of pro-inflammatory (*Nos2, Tgfb1, Il1β, Il6*) and fibrotic/wound healing associated genes (*Inhba, Spp1*, *Arg1, Pdgfb*, and *Fn1*) (**Figure 4C**). MHC II IMs express angiogenesis (*Col3a1* and *Pdgfb*), pro-inflammatory (*Ccl3* and *Mmp9*), and fibrotic markers (*Ccl24*). CellChat analysis revealed that Spp1 signaling from Mono/Macs towards neutrophils was elevated in infected lungs (**Supplemental Figure 3E**).

**Figure 4:**
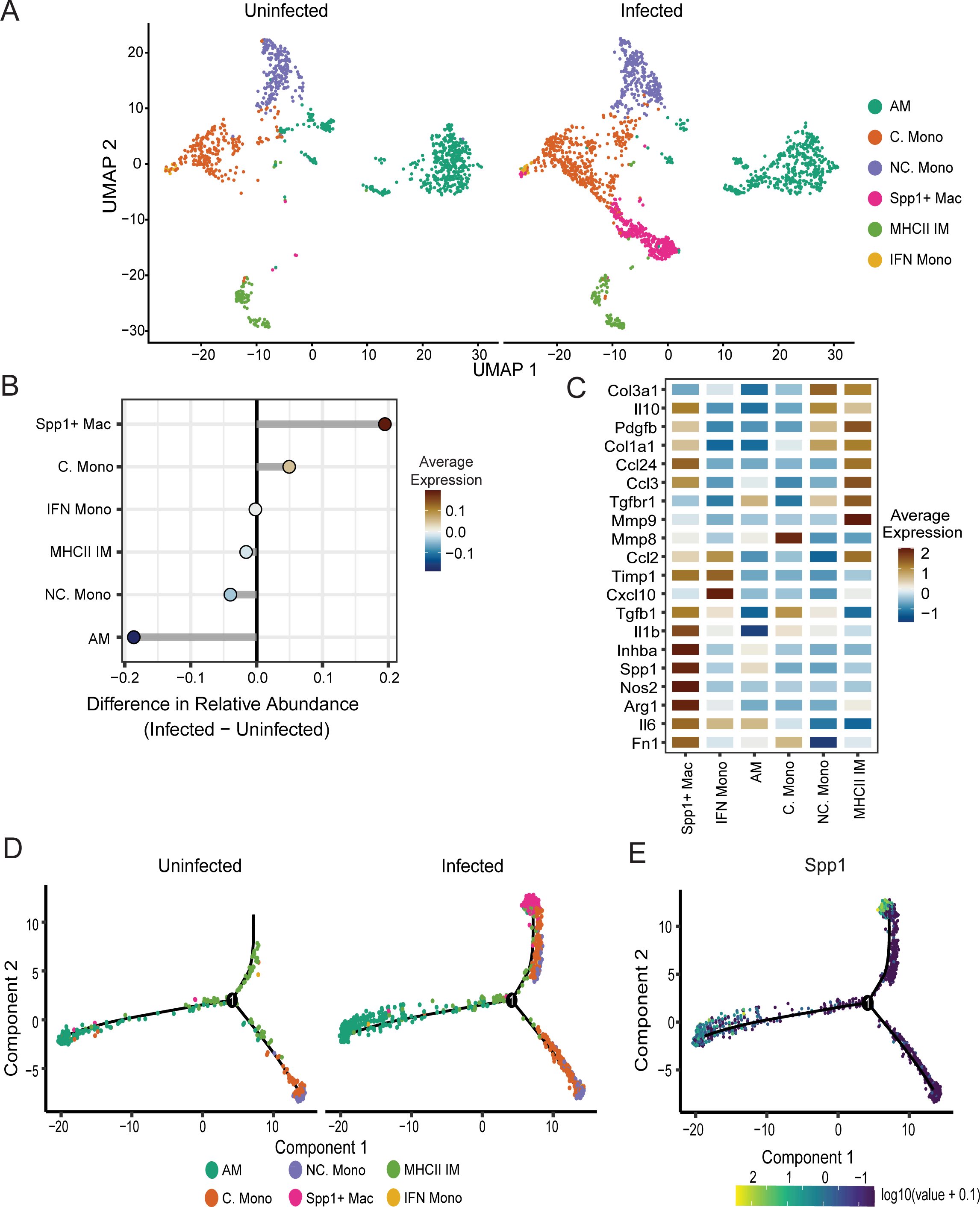
Spp1+ macrophages emerge following infection and display a fibrotic phenotype. (A) UMAP visualization of cell clusters identified in each experimental group. (B) Relative cellular proportions of monocyte/macrophage subsets. (C) Heat map showing expression level of select fibrotic and inflammatory signaling genes across monocyte and macrophage subclasses. Trajectory analysis on monocyte/macrophage subclusters, colored by identity (D) or Spp1 expression (E).

To understand macrophage polarization, trajectory analysis was applied to the Mono/Mac clusters on the combined uninfected and infected data, revealing three distinct lineages (**Figure 4D**, **S3C**). Lineage one consisted of classical and non-classical monocytes. Lineage two was comprised of alveolar macrophages, reflecting a distinct lineage from other subsets. Lineage three encompassed MHC II IM in uninfected samples and expanded to include monocyte populations and Spp1^+^ Macs in infected samples (**Figure 4D**). Pseudotime analysis indicated that classical and non-classical monocytes move towards the Spp1^+^ lineage after infection (**Figure S3C**), and *Spp1* expression is highest at the termination of lineage three (**Figure 4E**). Interestingly, moderate expression of *Spp1* was also observed at the termination of lineage two, suggesting that alveolar macrophages begin to express *Spp1* after infection. Overall, the gene expression profiles observed in the monocyte/macrophage subsets indicated notable shifts in differentiation states upon infection, with the striking emergence of an Spp1^+^ macrophage population that displays high expression of fibrotic genes.

### *Coccidioides* associate with neutrophils and Spp1^+^ macrophages *in vivo*

Immune responses against fungal infections involve direct interaction with fungi, activation of immune cells, and the release of paracrine signals to alert and recruit other immune cells to the site of infection. To probe the fungal-contact-dependency of the cellular responses in the lung, we employed fluorescently labeled fungal spores to separately assess responses of cells directly in contact with *Coccidioides* versus bystander cells. We adapted a fluorescent labeling scheme previously used to label *B. anthracis* spores with an Alexa Fluor 488 (AF488) amine-reactive dye (*27*). AF488 robustly and uniformly labeled arthroconidia without impacting fungal viability (**Figure S5A, B**). AF488 labeled *Coccidioides* were detected interacting with macrophages *in vitro* via fluorescence imaging and flow cytometry (**Figure S5C, D**). Flow cytometric analysis of lungs from mice infected with AF488 labeled *Coccidioides* revealed that the majority (>95%) of the cells interacting with *Coccidioides in vivo* were CD45^+^ immune cells (**Figure 5A, S5E**).

**Figure 5:**
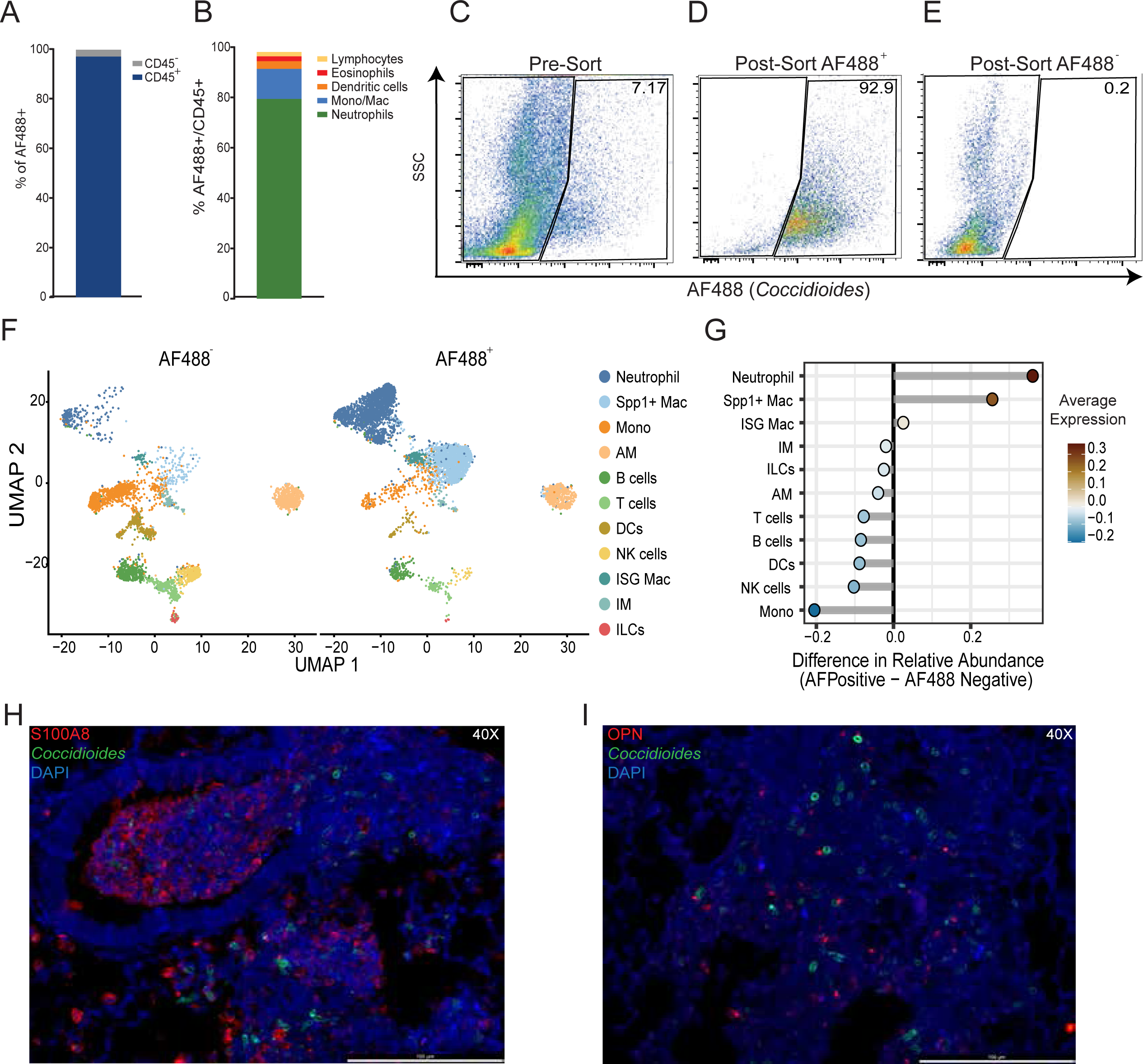
Neutrophils and macrophages associated with Coccidioides. Mice were infected intranasally with 1×106 AF488 labeled Coccidioides, and lungs harvested 24 hpi and processed for flow cytometric and scRNAseq analysis. Percentage of AF488+ cells isolated from the lungs of that are CD45+ and CD45– (A). Percentage of AF488+ cells corresponding to the indicated immune cell subtypes (B). Flow plots indicate the pre-sort (C) and post-sort purities of AF488+ (D) and AF488– (E) populations. UMAP visualization of cell clusters identified in each experimental group (F). Relative cellular proportions in AF488+ versus AF488– populations (G). Immunohistochemistry (IHC) of lung sections co-visualizing AF488-labeled Coccidioides with neutrophils (S100A8+) (H) or cells expressing OPN (I). Images were taken at 40X magnifications and counterstained with DAPI. n=3 uninfected and n=8 infected for flow cytometry. n=3 pooled per condition for sequencing. IHC n=3 per condition. Representative images are shown from a single animal.

∼80% of the total CD45^+^ AF488^+^ population were neutrophils (**Figure 5B, S5E-F**). The second most abundant cell type was monocytes/macrophages (AM, IM, and monocytes), collectively representing approximately 12% of the CD45^+^ AF488^+^ population. The remaining 8% were comprised of eosinophils, dendritic cells, and lymphocytes. The flow cytometry data was in line with scRNAseq results (**Figure 1C**) that demonstrated neutrophils and Mono/Mac cells were highly enriched in infected lungs and furthermore, were the most likely cell types to interact with fungal spores *in vivo* 24 hpi.

Next, we employed scRNAseq to elucidate gene expression changes upon contact with *Coccidioides*. Cells were harvested from lungs 24 hpi as before and enriched for immune cells by bead sorting for CD45^+^ cells. Cells were then sorted into AF488^−^ and AF488^+^ populations using fluorescence-activated cell sorting (FACS), resulting in post-sort purities of >99% and 92.9% for AF488^−^ and AF488^+^ populations (**Figure 5 C-E**). scRNAseq was then performed on AF488^−^ and AF488^+^ samples and unsupervised clustering of the data resulted in the identification of 11 cell clusters (**Figure 5F** and **S5G**). Consistent with flow cytometry data (**Figure 5B, S5E-F**), the relative abundance comparison between sequenced AF488^+^ and AF488^−^ populations indicated that neutrophils and Spp1^+^ macrophages were the most enriched cell types in the AF488^+^ sample (**Figure 5G**). The relative abundance of other immune cell types, such as B, T, NK, and dendritic cells, also aligned with flow cytometry data, confirming that these cell types are not frequently associated with *Coccidioides in vivo* 24 hpi (**Figure 5B, G**). Interestingly, monocytes were the cell population most highly enriched in the AF488^−^ sample. The corresponding high frequency of monocytes in the AF488^−^ sample and high frequency of Spp1^+^ macrophages in the AF488^+^ sample suggested that monocytes may be differentiating into Spp1^+^ macrophages upon contact with *Coccidioides.* This is further supported by the trajectory analysis in **Figure 4D**, which indicated that monocytes move towards the Spp1*^+^* lineage upon infection. Immunohistochemistry further confirmed the interaction of *Coccidioides* with neutrophils and Osteopontin (OPN)*^+^*cells *in vivo* (**Figure 5H, I**). AF488^+^ *Coccidioides* were visualized surrounded by neutrophils (S100A8*^+^*) (**Figure 5H**). OPN, the protein produced by *Spp1* gene, staining is observed intracellularly (purple) and extracellularly (red). OPN^+^ cells are seen surrounding *Coccidioides* and OPN^+^ localization on the cell surface is oriented towards *Coccidioides* (**Figures 5I**). Collectively, these data demonstrate that neutrophils and monocyte/macrophage cells associate with *Coccidioides* rapidly upon infection and suggest a novel mechanism by which monocytes differentiate into Spp1*^+^* cells upon contact with *Coccidioides*.

### *Spp1* expression is contact dependent

To verify that Spp1^+^ macrophage differentiation is contact dependent, macrophages were exposed to *Coccidioides in vitro* directly, or indirectly via conditioned media (cm), and expression of the gene signature was evaluated by RNA-seq. M-CSF and GM-CSF are both commonly used to induce macrophage differentiation from bone marrow *in vitro,* resulting in macrophages with distinct phenotypic characteristics (*28, 29*). We differentiated macrophages with each cytokine, then exposed them to *Coccidioides in vitro* to determine which cells would respond to fungal infection by upregulation of *Spp1.* Overnight incubation of GM-CSF differentiated bone marrow derived macrophages (GM-BMDM) with *Coccidioides* arthroconidia resulted in robust induction of *Spp1* expression (**Figure 6A**), whereas exposure of M-CSF differentiated bone marrow derived macrophages (M-BMDM) to *Coccidioides* resulted in significant cell death, as well as no induction of *Spp1* in surviving cells (**Figure 6A**).

**Figure 6:**
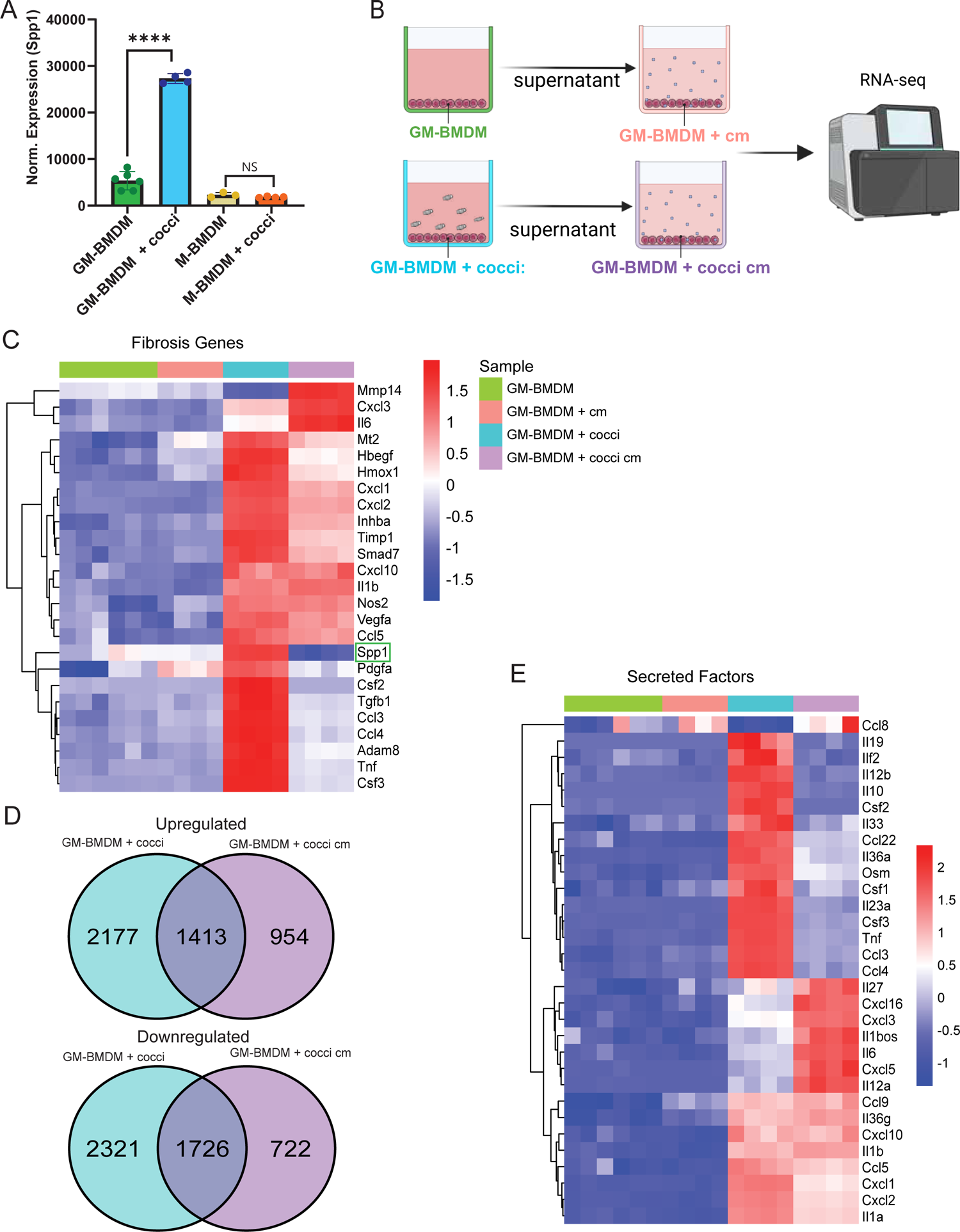
Spp1 expression is dependent on contact with Coccidioides. Bone marrow-derived macrophages (BMDM) were differentiated with GM-CSF (GM-BMDM) or M-CSF (M-BMDM) and exposed to 500,000 Coccidioides arthroconidia. RNA was isolated 24 hpi and RNA sequencing performed. Normalized expression of Spp1 (A). Schematic of conditioned media experiment (B). Heat map depicting expression of selected fibrosis-related genes (C). Genes differentially expressed between stimulated and unstimulated conditions (D). Heat map depicting expression of selected cytokine/secreted factor genes (E).

Next, to determine if *Spp1* induction was contact-dependent, fresh GM-BMDM were exposed to cm collected from control GM-BMDM or GM-BMDM incubated with *Coccidioides* (GM-BMDM + cocci) and RNA-seq was conducted on RNA isolated from GM-BMDM, GM-BMDM + cm, GM-BMDM + cocci, and GM-BMDM + cocci cm (**Figure 6B**). Fibrosis-related genes identified as highly expressed in Spp1^+^ macrophages *in vivo* (**Figure 4C**) were evaluated within each condition. Limited expression was observed in both control conditions, as expected (**Figure 6C**). Robust upregulation of fibrosis-related genes was observed in macrophages in contact with *Coccidioides,* including *Spp1, Tgfb1, Timp1, Adam8,* and *Inhba* (**Figure 6C**). While there was some overlap in gene expression patterns between cells in direct contact with *Coccidioides* and those exposed indirectly via cm, *Spp1* expression was absent in the GM-BMDM + cocci cm condition, and expression of most fibrotic genes was higher in the GM-BMDM + cocci vs GM-BMDM + cocci cm condition (**Figure 6C**), indicating that *Spp1* expression, as well as the expression of many fibrotic genes, was dependent upon contact with *Coccidioides*.

GM-BMDM + cocci cm initiated robust changes in gene expression, with 2367 and 2448 genes significantly up or downregulated, respectively (**Figure 6D**). Thus, we can infer that direct exposure of GM-BMDM to *Coccidioides* resulted in the release of secreted factors that activated GM-BMDM *in trans.* To probe the landscape of the secretome of GM-BMDM + cocci, we analyzed expression of a panel of secreted factor genes (**Figure 6E**). Contact with *Coccidioides* resulted in the upregulation of many pro-inflammatory cytokine genes, including *Il33*, *Il12b, Csf1*, *Csf2, Il23a*, *Tnf*, *Ccl3*, and *Ccl4*. Interestingly, while some overlap in cytokine gene expression was observed between conditions, including *Il1b, Cxcl10, Ccl5, Cxcl1,* and *Cxcl2,* distinct patterns of cytokine gene expression were observed in GM-BMDM + cocci vs GM-BMDM + cocci cm conditions. Notably, robust *Il6* expression was observed only within the GM-BMDM + cocci cm condition. Taken together, these data confirm that direct contact with *Coccidioides* is necessary for the induction of *Spp1* expression in macrophages, and furthermore, this contact promotes robust expression of fibrosis-related genes as well as a panel of cytokines that in turn robustly stimulate a distinct activated phenotype *in trans*.

## Discussion

The lung microenvironment presents a complex and dynamic space where the immune response to *Coccidioides* infection unfolds. This response is orchestrated by a range of innate immune cells and is influenced by the unique interactions between the pathogen and host that have not been fully characterized at the cellular and molecular level. Using a comprehensive transcriptomics approach, we have dissected the immediate cellular dynamics triggered by a high dose of *Coccidioides* and identified pivotal cellular players initiating the antifungal immune response. The immune response to high dose attenuated *C. posadasii* Δ*cts2*/Δ*ard1*/Δ*cts3* 24 hpi is characterized by the rapid recruitment and activation of neutrophils and OPN^+^ cells that aggregate around *Coccidioides* and differentiate into PD-L1^+^ neutrophils and Spp1^+^ macrophages. Chemotaxis and proliferation gene signatures suggest innate immune populations are recruited to the lung and actively differentiate in response to localized signaling. This early immune activation is crucial for controlling the initial fungal invasion and setting the stage for subsequent adaptive immune responses but may also drive inappropriate inflammation, damage, and fibrosis.

Surprisingly few changes occurred among the non-immune cells within the lung during infection, although these cells are critical for maintaining homeostasis and barrier function during infection. Within this lung subset, an increase in AT2 epithelial cells relative to other non-immune cell types was observed. The main functions of AT2 cells are production and secretion of surfactant and as progenitors that replenish epithelial cells after lung injury (*30*). Expansion of this population suggests an attempt to prevent fungal lung entry using surfactant, or damage to the epithelium by *Coccidioides* entry and growth. Avirulent *Coccidioides* is rounded and enlarged within 24 hpi in the lungs while transitioning from arthroconidia to the parasitic form (*31*), and the transition from arthroconidia to spherules likely causes tissue damage as the spherules become very large within the tissue. A recent scRNAseq study examining primary human airway epithelial cells infected with *Coccidioides* reported hypoxic responses upregulated during infection (Harding et al., 2024, submitted for publication). Notably, hypoxia is known to contribute to the development of fibrotic conditions within the lung (*32–34*). Fibroblasts exhibited enrichment in fibrosis-related pathways, which may suggest crosstalk with pro-fibrotic macrophages that emerge after infection. The molecular interactions suggest a combined repair and protective response by the barrier cells within the lung following fungal infection, and highlight the complexity of the lung environment, but also suggest poor barrier activation against this fungus which may be shifted towards a pro-fibrotic response.

Unsurprisingly, neutrophils were rapidly recruited to the lung and constituted the largest cellular increase following *Coccidioides* infection. Neutrophils engulf and kill immature spherules, such as those that form with the attenuated strain used in this study, and are required for vaccine efficacy against *Coccidioides* (*35*). Neutrophils were subclustered into five subsets, with largely overlapping gene signatures, likely indicating active differentiation within the lung microenvironment 24 hpi. The presence of Circulating and Early neutrophils undergoing differentiation suggests active granulopoiesis in the bone marrow, seeding the lung tissue. Trajectory analysis indicates that Circulating and Early neutrophils differentiate into PD-L1^+^ neutrophils, a population known to negatively regulate antifungal activity (*36–38*). During *Candida* infections, PD-L1^+^ neutrophils inhibit antifungal activity through autocrine signaling involving CXCL1 and CXCL2. This signaling is associated with neutrophil migration and, in the context of *Candida* infections, leads to their movement to and retention within the bone marrow thereby, inhibiting an effective neutrophil response. PD-L1^+^ neutrophils exacerbate acute respiratory distress injury via the PI3K/Akt pathway by enhancing autophagy and inflammation (*39*). Further studies are needed to define if and how PD-L1^+^ neutrophils contribute to a maladaptive response during *Coccidioides* infection, and to determine if PD-L1^+^ neutrophils arise through fungal contact-dependent mechanisms, similar to Spp1^+^ macrophage differentiation. The mechanisms driving PD-L1^+^ neutrophil differentiation and activation are also unknown, however this population also dominates the lung during virulent *Coccidioides* infection, although at later stages post-infection (Davalos et al., 2024, submitted for publication). Differences in the differentiation kinetics of PD-L1^+^ neutrophils between these studies may be due to fungal dosing or fungal evasion mechanisms that require spherule maturation and may explain why neutrophils are required for development of vaccine immunity to *Coccidioides*, while neutrophil depletion does not influence disease mortality (*35*)(*35*)(*34*)(*33*)(*32*)(*32*).

The second most prominent cellular response observed in our study is the monocyte-derived macrophages that rapidly differentiate within the tissue by upregulating *Spp1*. *Spp1* expression is an emerging biomarker in various diseases, with the presence of Spp1^+^ macrophages being linked to poor prognosis in severe COVID-19, active TB infection, lung fibrotic diseases, and cancer (*40–49*). Trajectory analysis indicated classical monocytes transition into Spp1^+^ macrophages following *Coccidioides* infection, and *in vitro* experiments indicated this differentiation is dependent on fungal contact. OPN, produced by the *Spp1* gene, is a versatile secreted molecule, inducing pro- and anti-inflammatory responses through several pathways, including the Dectin-1 receptor which is crucial for recognizing *Coccidioides* (*50–52*). Future studies are necessary to define the surface molecules and downstream signaling pathways expressed by monocytes that engage with *Coccidioides* and result in Spp1^+^ macrophage differentiation. In a mouse model of *Candida* infection, *Spp1* deficiency completely ameliorates disease lethality (*53*). In severe COVID-19, elevated OPN plasma levels activated CD14 monocytes and PD-L1^+^ neutrophils, contributing to disease progression (*50*). Together these findings suggest that upregulation of *Spp1* may initially represent an attempt to control disease following direct contact with *Coccidioides* but may ultimately be detrimental to infection outcomes. Thus, Spp1^+^ macrophage polarization in *Coccidioides* may represent the beginning of a maladaptive immune response at 24 hpi that promotes PD-L1^+^ neutrophil differentiation (*54, 55*). Additional studies are needed to clarify the role of Spp1^+^ macrophages in disease outcomes; however, Spp1^+^ macrophages also develop in response to virulent *Coccidioides* infection (Davalos et al., 2024, submitted for publication), expanding from 5 to 14 days post infection and demonstrating a similar fibrotic phenotype.

Our comprehensive analysis reveals that the early immune response to *Coccidioides* infection involves a rapid influx and activation of neutrophils, particularly PD-L1 expressing subsets, and the differentiation of monocytes into Spp1^+^ macrophages. These findings highlight the dynamic nature of the lung microenvironment during the initial stages of infection and underscore the complex interplay between pathogen clearance and tissue repair mechanisms. The emergence of Spp1^+^ macrophages, which exhibit both pro-inflammatory and fibrotic gene signatures, suggests a potential for maladaptive immune responses that could complicate long-term disease resolution. Understanding these early immune dynamics is crucial for developing targeted therapies to balance effective pathogen clearance with minimizing tissue damage, thereby improving clinical outcomes for Valley fever and other fungal infections exacerbated by climate change. Future studies should focus on the functional roles of these immune subsets in chronic infection stages and their impact on disease progression. Investigations could include examining the molecular mechanisms driving PD-L1^+^ neutrophil differentiation and activation, determining the specific contributions of Spp1^+^ macrophages to tissue fibrosis, and exploring potential therapeutic interventions to modulate these immune responses. Additionally, studies utilizing virulent strains and varying dosages could provide further insights into the kinetics and severity of immune responses, ultimately informing better strategies for prevention and treatment.

## Methods

### Fungal culture

*Coccidioides posadasii* (Δ*cts2*/Δ*ard1*/Δ*cts3*) derived from patient isolate C735 was used for all infections (NR-166 BEI Resources, Manassas, VA, USA). Liquid 2X Glucose 1X Yeast Extract (2X GYE) media (Fisher Scientific, Hampton, NH, USA) with 50 mg/mL hygromycin B (Invitrogen CAT# 10687010) was used to grow *Coccidioides* mycelia in a shaking incubator at 37°C for a week. Mycelia was then streaked onto solid 2X GYE agar and desiccated in an incubator at 30°C for a month to disassociate mycelia into individual arthroconidia. To harvest arthroconidia, the white fuzzy growth (arthroconidia) was scraped off with 1X PBS (VWR) and filtered through a 100 µM mesh filter. *Coccidioides* was then vortexed for 1 min to dissociate the arthroconidia and centrifuged at 12000 x *g* for 8 mins with break off at room temperature. The pellet was washed and vortexed with 30 mL of PBS and centrifuged again. After the second spin, 10 mL 1X PBS was added to the pellet and working stock stored at 4°C. Fresh arthroconidia stocks were prepared monthly. See Mead et al. for detailed protocol (*56*).

### Arthroconidia fluorescent labeling

Alexa fluor 488 (AF488, ThermoFisher, Cat #: A20000) was used to label arthroconidia spores as previously described (*27*). Briefly, AF488 was resuspended in 100 µl DMSO for the final concentration of 10 mg/mL, aliquoted and stored at −20°C. To label *Coccidioides*, 1 mL of fresh 1M NaHCO_3_ and 5 µl AF488 were combined in a 15 mL tube along with 6×10^6^ arthroconidia and 1X PBS added a final volume of 10 mL. The mixture was incubated at room temperature for 3 hrs, then centrifuged at 12000 x *g* for 8 min with break off at room temperature. Pellet was then washed with 15 mL PBS and centrifuged again, then resuspended to a final concentration of 1-10×10^6^ cells/40 µl).

### Mouse infections

6–8-week C57BL/6 female mice (JAX #000664, The Jackson Laboratories, Bar Harbor, ME, USA) were used for all animal experiments. Animal work was approved by the Lawrence Livermore National Laboratory Institutional Animal Care and Use Committee under protocol #309. Animals were housed in an Association for Assessment and Accreditation of Laboratory Animal Care (AAALAC)-accredited facility. Mice were infected intranasally with 1×10^6^ arthroconidia (with AF488 label or unlabeled) in 40 µl (20 µl per nostril) 1X PBS while under anesthesia (4-5% isoflurane (Covetrus) in 100% oxygen). For tissue harvest, animals were anesthetized under isoflurane and the whole animal was perfused with 20 mL sterile PBS containing 50,000 U/L sodium heparin (Sigma) *via* the left ventricle.

### Lung isolation and preparation of single cell suspensions

Following euthanasia and perfusion, lungs were removed and placed in a 100 mm petri dish containing 5 mL of digestion buffer (DMEM/F12 (Thermo Fisher) + 100 µg/mL DNase I (Roche cat# 11284932001) and 3 mg/mL collagenase 1 (Worthington cat# LS004197)) and minced with scissors into pieces approximately 1 mm in diameter. Minced pieces were transferred to a 15 mL conical tube and petri dish rinsed with another 5 mL digestion buffer, which was then added to the 15 mL tube. Lung homogenate was incubated for 30 mins in a 37°C shaking incubator at 150 rpm, with cells triturated with a 10 mL pipet at 15 min. At 30 min, supernatant was removed and transferred to a 50 mL conical tube and placed on ice. 10 mL of fresh digestion buffer was then added to remaining lung homogenate and incubated for another 30 min with shaking as above. Following the second 30 min incubation, remaining unhomogenized tissue was pipetted into a 70 µM mesh filter and manually dissociated with the flat end of a pellet pestle. Equal volume DMEM/F12 + 10% fetal bovine serum (FBS, Thermo Fisher) was added to cell suspension to neutralize digestion enzymes and pelleted by centrifugation at 500 x *g* at 4°C for 8 minutes. Red blood cells were removed by incubation in ACK lysis buffer (Thermo Fisher) for 5 min at room temperature. Cells were then resuspended DMEM/F12 + 10% FBS and counted on a Countess II (Thermo Fisher) automated cell counter prior to downstream analysis.

### Preparation of cells for scRNAseq

For whole lung sequencing, 3×10^6^ cells were fixed per mouse using 10x Genomics Fixation Buffer according to manufacturer’s instructions and fixed cells stored at −80°C prior to sequencing. When thawed, fixed cells were counted again using a Countess 3 (Thermo Fisher) and 500,000 cells each from 3 mice were pooled per reaction. For sequencing of sorted cells, following generation of single cell suspensions, immune cells were enriched using LS columns (Miltenyi Biotec) for MACS separation using anti-CD45 magnetic microbeads (Miltenyi Biotec, 130-052-301) according to manufacturer’s instructions, followed by resuspension in 10x Genomics Fixation Buffer and fixed overnight at 4°C. Fixed cells were pooled and sorted using a BD FACSAria Fusion and BD FACSMelody Cell Sorter to collect AF488^−^ or AF488^+^ populations. BD FACSDiva Software Version 8.0.1 and BD FACSChorus Software Version 2.0 Application Data Version 1.1.20.0. was used during acquisition to define the sorted populations. Sorted cell suspensions were resuspended in 10x Genomics Quenching Buffer and glycerol according to manufacturer’s instructions and stored at −80°C prior to sequencing.

### Single cell RNA sequencing (scRNAseq) Library Preparation

Single-cell RNA sequencing was performed using the 10x Genomics Chromium Single Cell Fixed RNA Profiling (Flex) Kit (cat# 100475), following the manufacturer’s protocol. Following the generation of single-cell suspensions as described above, 3×10^6^ cells per lung sample were fixed using 10x Genomics fixation buffers supplemented with paraformaldehyde to a final working solution of 4% according to manufacturer’s instructions and then stored at −80°C 10x Genomics long-term storage solution prior to library preparation. Each reaction was inclusive of 3 mice validated for fungal infection by plaque assay and pooled at 500,000 cells per mouse. Probe hybridization was then performed overnight followed by pooling before loading onto the Chromium X Controller (10x Genomics) to generate single-cell gel bead-in-emulsions (GEMs). Reverse transcription, cDNA amplification, and library amplification were performed according to the manufacturer’s recommendation. Quality control measures, including quantification and size distribution analysis, were performed using an Agilent Tape Station. Sequencing libraries were then sequenced on an Illumina NextSeq 2000 platform, producing paired end reads. Data processing, including alignment, barcode assignment, and gene expression quantification, was performed using the Cell Ranger software pipeline (v 7.2.0) from 10x Genomics (*57*). Alignments were carried out using the reference genome package refdata-gex-mm10-2020-A for GRCm38 (mm10), provided by 10x Genomics. Additionally, the “Chromium_Mouse_Transcriptome_Probe_Set_v1.0.1_mm10_2020-A” probe set from 10x Genomics was utilized.

### Flow cytometry

Following the generation of single-cell suspensions as described above, 1 million cells per condition were incubated with the following antibodies: CD11c PE Cy7 (BD, CAT# 558079, clone HL3, 1:500), SiglecF PE (Invitrogen, CAT# 12-1702-82, clone 1RNM44N, 1:500), CD11b BV 510 (Biolegend, CAT# 101263, clone M1/70, 1:500), Ly6C APC (#BD, CAT# 560595, clone AL-21, 1:500), Ly6G BV711 (BD, CAT# 563979, clone 1A8, 1:500). Samples were incubated with antibody cocktail for 30 mins at 4°C, washed, fixed with 4% paraformaldehyde for 1 h, then washed and resuspended in 200 µL of PBS for flow cytometry. Samples were acquired using the BD FACSAria™ Fusion Flow Cytometer utilizing the BD FACSDiva™ Software Version 8.0.1. Flow cytometry results were analyzed using FlowJo™ v10.10 Software (BD Life Sciences). All immune cell types were first gated on CD45 and further identified by expression of the following markers: lymphocytes: CD11b^−^CD11c^−^, alveolar macrophages (AM): CD11b^lo^SiglecF^+^, eosinophils: CD11b^hi^SiglecF^+^, neutrophils: CD11b^+^Siglec F^-^Ly6G^+^, dendritic cells (DCs): CD11c^+^MHC II^+^, monocytes: CD11b^+^SiglecF^-^Ly6G^-^ MHC II^−^Ly6C^+^, interstitial macrophages (IM): CD11b^+^SiglecF^−^Ly6G^−^MHC II^-^Ly6C^−^. Sequential gating strategy is visualized in **Figure S5E**.

### Immunohistochemistry (IHC)

Lungs for immunohistochemical staining were collected from infected mice at 24 hpi. After euthanasia and perfusion, the lungs were inflated and immersed in 10% neutral buffered formalin (NBF) at 4°C for 3 days with mild agitation. Lung samples were then paraffin-embedded and sectioned into 5 μM thick sections using a Leica RM2255 Microtome. The sections were deparaffinized using xylene and hydrated in a series of alcohol solutions. Antigen retrieval was performed at 65°C with Tris-EDTA (Abcam, ab93684) and non-specific sites were blocked with CAS-block (008120, Thermo Fisher Scientific). The sections were incubated with primary antibodies S100A8 (Abcam, ab92331) and OPN (AF808, R&D), followed by secondary antibodies (A11037 and A11058, Thermo Fisher Scientific). Finally, the slides were mounted with ProLong Gold Antifade Mounting with DAPI (ThermoFisher, CAT# P36935) and imaged using a Leica DM4 B microscope with 20X and 40X objective.

### scRNAseq data analysis

#### Full data set analysis

Analysis was initially performed using Seurat (v4.3.0) (*58*)and R (v4.3.2) but transitioned to Seurat (v5.1.0) (*59*)and R (v4.4.0). Seurat objects were created for each sample using the CreateSeuratObject function with default parameters and subsequently merged into a single object. For the analysis, we utilized uninfected lung sample data from the D0 timepoint, as described in Davalos et al. (2024). Quality control of the datasets involved retaining cells with at least 500 counts, cells with at least 300 expressed genes, and cells with less than 10% mitochondrial genes (nCount_RNA ≥ 500, nFeature_RNA ≥ 300, and percent_mt < 10). We excluded lowly expressed genes by retaining only those genes that were expressed in a minimum of 10 cells.

### Broad analysis of the entire dataset

Data was normalized using ‘NormalizeDat’ function using default parameters. Highly variable genes (HVGs) were identified using the ‘FindVariableGenes’ function. Before dimension reduction, the data was scaled with the effects of cell counts and mitochondrial percentages regressed out using ‘ScaleDat’ (vars.to.regress=c(“percent_mt”, “nCount_RNA”). After performing principal component analysis (PCA) using thèRunPCÀ function, the top 50 principal components were selected for downstream integration, clustering, and dimension reduction.

Data integration was performed using Harmony (v1.2.0) (*60*) with the grouping variable set to “orig.ident” which contained both samples. Clustering analysis was performed using FindNeighbors and FindClusters, with the reduction parameter set to “harmony” and resolution of 0.4. Subsequently, a non-linear dimensionality reduction was conducted using uniform manifold approximation and projection (UMAP), with ‘RunUMAP’ (reduction = “harmony”, umap.method = “uwot”).

To distinguish different cell populations, cell annotation utilizing differential expression, known marker genes, and a scRNA-seq resource LungMAP (*61*) were used. Labels were transferred (lineage_level1, lineage_level2, celltype_level2) from the mouse LungMAP scRNAseq object (*36*) by using two functions ‘FindTransferAnchors’ (reference = lungmap, query=ourdata), and ‘TransferDat’ (dims = 1:30). Next, ‘FindAllMarkers’ (only.pos = true) was used to identify cluster marker genes and annotated cell clusters using known marker genes. The predicted labels and the label scores from LungMAP were overlayed to confirm appropriate cell labeling across all populations.

Lastly, the relative abundance of cell types was calculated by determining the proportion of each cell type within each sample. The difference in proportions between two samples was then computed for each cell type. These differences were used to visualize changes in cell type abundance between the samples.

### Neutrophil subset

To analyze neutrophils and their various subsets, we followed the same approach used for the entire dataset with a few modifications. Given the smaller size of the neutrophil subset we identified 2000 HVGs. For dimension reduction, integration, and clustering 10 principal components and a cluster resolution of 0.5 was used. For marker gene identification the same method was employed as described above in 2.10b. To identify biologically relevant pathways, the top 100 differentially expressed genes (DEGs) that met our filtering criteria (p-value adjusted < 0.05 and average log2 fold change ≥ 1) were selected. To perform enrichment analysis, clusterProfiler (v4.12.0) (*62*)‘compareCluster’ function with biological process ontology (ont = BP) were utilized. To perform a trajectory analysis, we used Slingshot (v2.12.0) (*26*) ‘slingshot’ (extend = n, stretch = 0, start.clus = Circulating Neu.) function on the UMAP embedding. Lastly, we utilized functions from the Single-Cell Pipeline (SCP) (v0.5.6) package to fit Generalized Additive Models (GAMs) for analyzing gene expression along pseudotime in scRNAseq data (Pedersen et al., 2019). Briefly, a GAM was applied to each gene, with gene expression as the dependent variable and pseudotime as the independent variable, incorporating smooth functions to capture non-linear relationships. This approach enabled the detection of changes in gene expression over pseudotime. We generated a heatmap of the dynamic features along the pseudotime trajectory using functions ‘RunDynamicFeatures’(n_candidates = 2000, seed = 2024, minfreq=0), ‘DynamicHeatmap’(use_fitted = TRUE, n_split = 3, r.sq = 0.09, dev.expl = 0.1, num_intersections = NULL).

### Myeloid subset

To further analyze myeloid cells and their various subsets, we followed the same approach used for the entire dataset (2.10b) with a few modifications. Given the smaller size of the myeloid subset we identified 2000 HVGs. For dimension reduction, integration, and clustering we used 30 principal components and a cluster resolution of 0.8. For marker gene identification we employed the same method as described above. Calculation of relative abundance was performed as stated above. Trajectory analysis was performed using Monocle2 (v2.32.0) (*63*) with the following sequence of functions and parameter adjustments: ‘ as.CellDataSet’, ‘estimateSizeFactors’, ‘estimateDispersions’, ‘reduceDimension’ (reduction_method = “DDRTree”, pseudo_expr = 1), ‘orderCells’(reverse=FALSE).

### Sorted AF488^-^ and AF488^+^ cells

For the analysis of the sorted data, the same approaches were used as in 2.10b with the following modifications. Seurat objects for each sample using CreateSeuratObject with default settings were constructed and merged into a unified object. To ensure data quality, we implemented several filters: retaining cells with at least 200 and less than 5000 counts, at least 200 and less than 5000 expressed genes, and less than 10% mitochondrial genes (200 >= nCount_RNA <= 5000, 200 >= nFeature_RNA <= 5000, and percent_mt < 10). Again, our analysis was restricted to genes that expressed a minimum of 10 cells. We identified 3000 HVGs. For dimension reduction, integration, and clustering 40 principal components and a cluster resolution of 0.4 were used. Annotation and calculation of relative abundance were performed as stated above.

### *In vitro* macrophage differentiation and infection

Cells were isolated from bone marrow as previously described (*64*). After processing to single cell suspension, red blood cells were lysed, and cells counted as described above. 500,000 cells per well were plated in a 12-well tissue culture treated plate in 2 mL DMEM/F12 + 10% FBS supplemented with 100 ng/mL M-CSF (Thermo Fisher, cat# PMC2044) or 100 ng/mL GM-CSF (R&D systems cat# 415-ML) to generate M-BMDM or GM-BMDM, respectively. Macrophages were differentiated for 7 days *in vitro,* with 1 mL media supplementation after 3 days in culture and a half media change on day 5. For direct contact experiments, 500,000 arthroconidia were added to macrophages in DMEM/F12 + 10% FBS and incubated for 24 h. Supernatants were collected, centrifuged at 2,000 x *g* for 10 minutes to remove cellular debris and filtered through a 0.22 µM filter to generate conditioned media. Conditioned media was also collected from mock-infected cells. For conditioned media experiments, GM-BMDM were incubated for 24 h with conditioned media supplemented with an additional 5% FBS from GM-BMDM infected with *Coccidioides* or mock-infected cells. RNA was isolated from all conditions using RNeasy RNA mini kits (Qiagen, cat #74014) according to manufacturer’s instructions.

### RNA-Sequencing library preparation and analysis

Sequencing library preparation was performed using Illumina Stranded mRNA Prep, Ligation (Illumina, cat # 20040534) according to manufacturer’s recommendations. Sequencing was performed on an Illumina NextSeq 2000 using P2 reagents. The raw RNA sequencing data was aligned to the mouse genome (mm10) using STAR (*65*), and mapped reads were counted using featureCounts (*66*). Subsequently, data normalization was performed using the Trimmed Mean of M-values (TMM) method (Robinson and Oshlack, 2010). Differentially expressed (DE) genes were identified using edgeR (version 4.2.1) (*67*) in R (version 4.4.1). Genes were considered significantly expressed when its false discovery rate (FDR) adjusted p-value was <0.05 and fold change was 1.5. Pathway and biological process enrichment analyses were conducted using the ToppGene Suite (*68*), based on the DE genes identified. Heatmaps were generated using the ‘pheatmap’ package (version 1.0.12) in R.

### Statistics

Data was evaluated using one- and two-way ANOVA followed by either Tukey’s or Šídák’s multiple comparisons test. Grubbs’ outlier calculations were performed as needed for flow cytometry data. GraphPad Prism version 10.2.2 for Windows was used, GraphPad Software (Boston, Massachusetts USA).

## Data Availability

The single-cell RNA sequencing and bulk RNA sequencing data have been deposisted in the NCBI Gene Expression Omnibus (GEO). GEO accension numbers will be provided upon acceptance of the manuscript.

## Acknowledgements and Funding

Funding for this research was provided by internal Lawrence Livermore National Laboratory Directed Research and Development funds (22-ERD-010 to D.R.W. and G.G.L.). N.M. was supported by UC-National Lab In-Residence Graduate Fellowship (L22GF4531). The funders had no role in study design, data collection and analysis, decision to publish, or preparation of the manuscript. This work was performed under the auspices of the U.S. Department of Energy by Lawrence Livermore National Security, LLC, Lawrence Livermore National Laboratory under Contract DE-AC52-07NA27344.

## Author contributions

A. N. Miranda: Conceptualization, Investigation, Methodology, Visualization, Funding acquisition Writing—original draft.

O.A. Davalos: Conceptualization, Investigation, Methodology, Formal Analysis, Visualization, Data Curation.

A. Sebastian: Conceptualization, Investigation, Methodology, Formal Analysis, Visualization, Data Curation, Resources.

M.V. Rangel: Investigation, Visualization, Data Curation.

N.F. Leon: Investigation, Visualization, Data Curation.

B.M. Gorman: Methodology, Visualization, Data Curation.

D.K. Murugesh: Investigation, Data Curation.

N.R. Hum: Conceptualization, Investigation, Methodology, Data Curation, Resources.

G.G. Loots: Funding acquisition.

K.K. Hoyer: Conceptualization, Writing—original draft, Supervision, Funding Acquisition.

D.R. Weilhammer: Conceptualization, Investigation, Project administration, Methodology, Visualization, Resources, Funding acquisition, Supervision, Writing—original draft.

All authors contributed to review and editing.

Disclosures: The authors declare no competing interests exist.

## Abbreviations

AF488: Alexa Fluor 488
AM: Alveolar Macrophage
AT2: Type 2 Pneumocytes
BMDM: bone marrow derived macrophages
C: cluster
CD: cluster of differentiation
cm: conditioned media
C Mono: Classical Monocyte
cocci: *Coccidioides*
C. posadasii: Coccidioides posadasii
Cd274: CD274 Antigen (PD-L1)
Circulating Neu: Circulating Neutrophils
COVID-19: Coronavirus Disease 2019
CXCL: Chemokine (C-X-C motif) Ligand
Cxcl2: Chemokine (C-X-C motif) Ligand 2
DCs: Dendritic Cells
DEGs: differentially expressed genes
hpi: hours post infection
Early Neu: Early Neutrophils
FACS: fluorescence-activated cell sorting
GM-CSF: Granulocyte-macrophage colony-stimulating factor or *CSF2*
GO: gene ontology
HVGs: Highly variable genes
IFN-γ: interferon gamma
IL-6: Interleukin 6
ILCs: Innate Lymphoid Cells
ISG: Interferon-Stimulated Gene
ISG Mono: Interferon-Stimulated Gene-expressing Monocyte
ISG Neu: ISG-expressing Neutrophils
Mac: macrophage
MHCII IM: Major Histocompatibility Complex Class II Interstitial Macrophage
Mono/Mac: Monocytes and Macrophages
neu: neutrophil
NC Mono: Non-Classical Monocyte
OPN: Osteopontin (Spp1)
PI3K/Akt: phosphoinositide 3-kinase (PI3K)/protein kinase B (AKT) pathway
PC: Principal Component
PCA: Principal Component Analysis
PD-L1: Programmed Death-Ligand 1
PD-L1+ Neu: PD-L1+ Neutrophils
Prolif.: Proliferating cells
RNA-seq: RNA sequencing
S100a8: S100 Calcium Binding Protein A8
scRNAseq: single-cell RNA sequencing
Spp1: Secreted Phosphoprotein 1 (OPN)
T/NK: T and Natural killer cells
Th1: T helper 1
Th17: T helper 17
TNFA or TNF-α: Tumor Necrosis Factor Alpha
Tregs: Regulatory T cells
UMAP: Uniform Manifold Approximation and Projection

## Supplemental figure legends

**Supplemental Figure 1:**
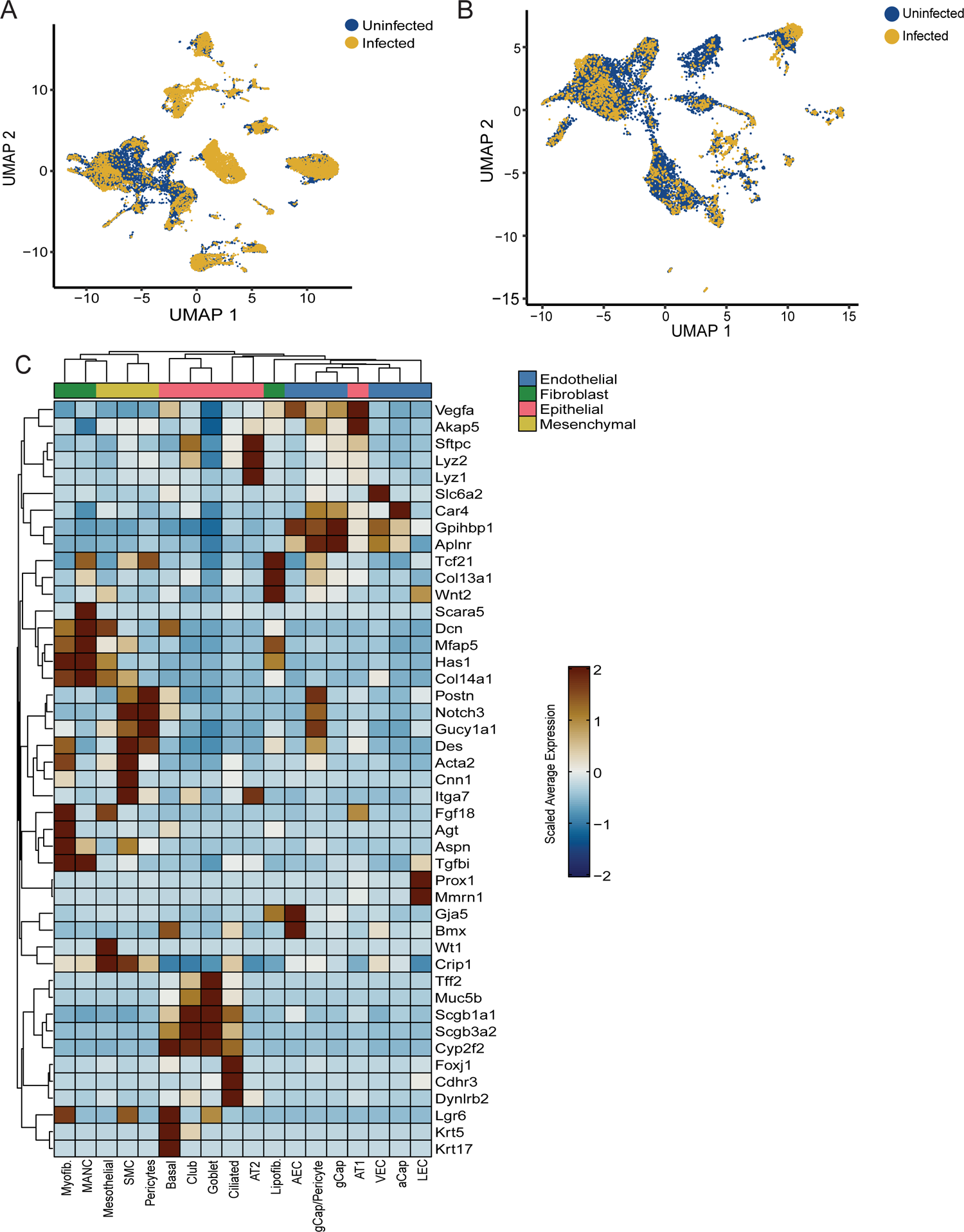
UMAPs and heat map of immune and non-immune cells. (A) UMAP overlay of all lung cells in the infected and uninfected samples from Figure 1B. (B) UMAP overlay of non-immune lung cells in the infected and uninfected samples from Figure 2A-B. (C) Heatmap of non-immune cells classification and gene signatures from Figure 2.

**Supplemental Figure 2:**
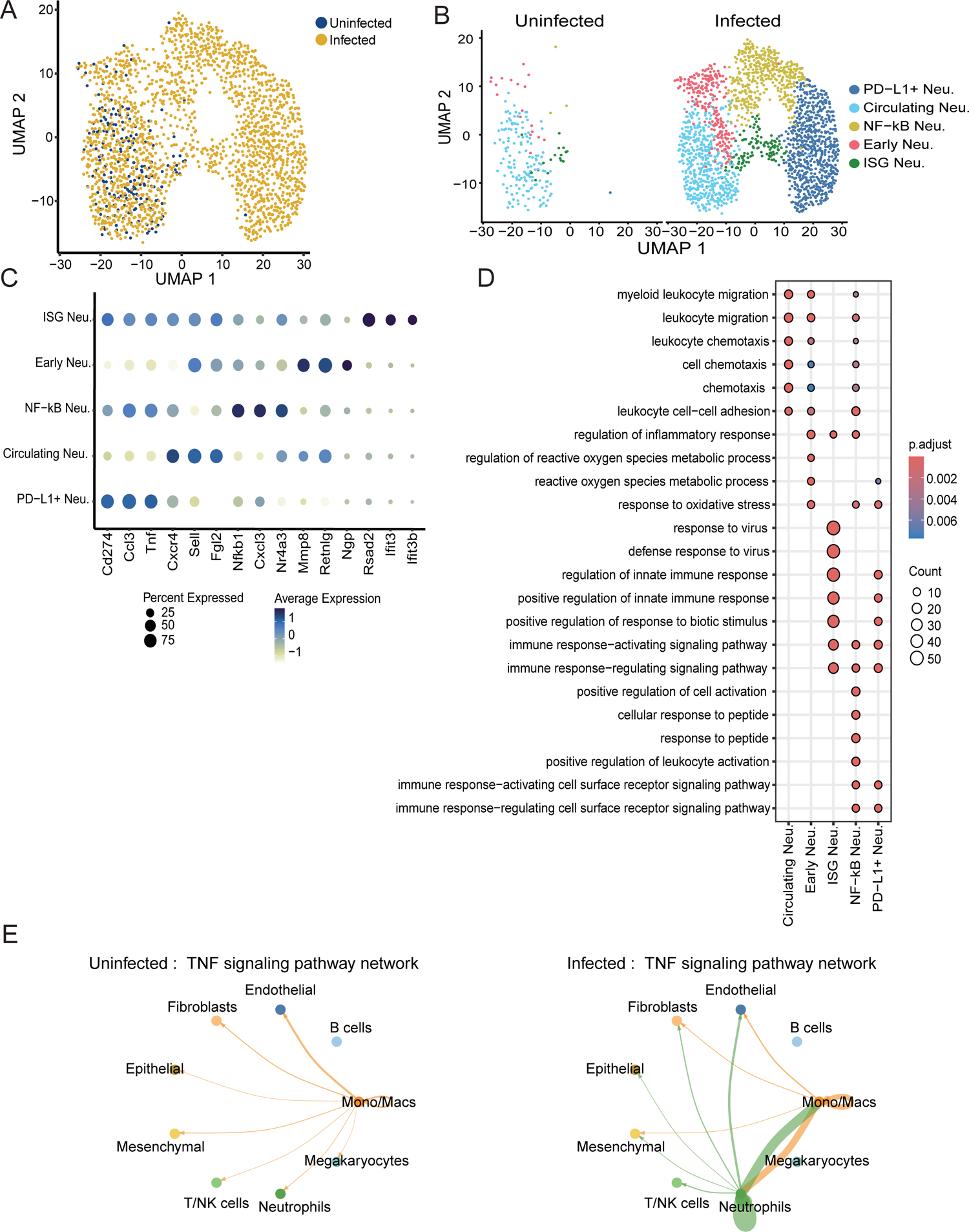
Extra neutrophil data. (A) Subclasses of neutrophils overlayed between uninfected and infected. (B) UMAP of uninfected and infected neutrophil subclasses. (C) Dot plot demonstrating top three gene signatures of each neutrophil subclass. (D) GO showcasing increased and decreased pathways associated with each neutrophil subclass. (E) CellChat identified TNF signaling in uninfected and infected samples.

**Supplemental Figure 3:**
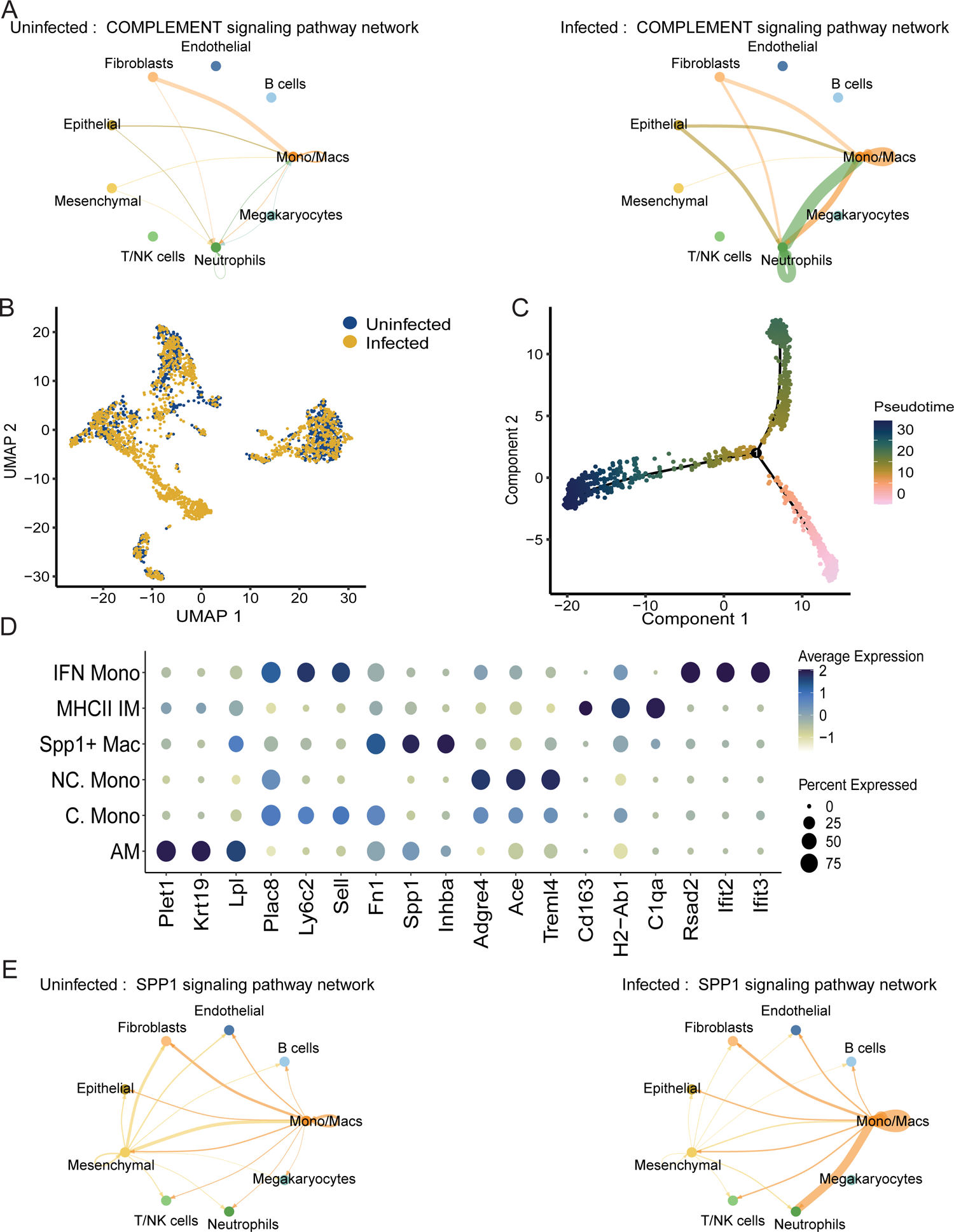
Extra macrophage data. (A) Complement signaling is through CellChat in uninfected and infected samples. (B) UMAP overlay macrophage subclusters from the uninfected and infected samples. (C) Pseudotime analysis on the macrophage subclusters for trajectory analysis. (D) Dot plot showing the top 3 genes expressed by each macrophage subclass. (E) Spp1 signaling via CellChat in infected and uninfected samples.

**Supplemental Figure 4:**
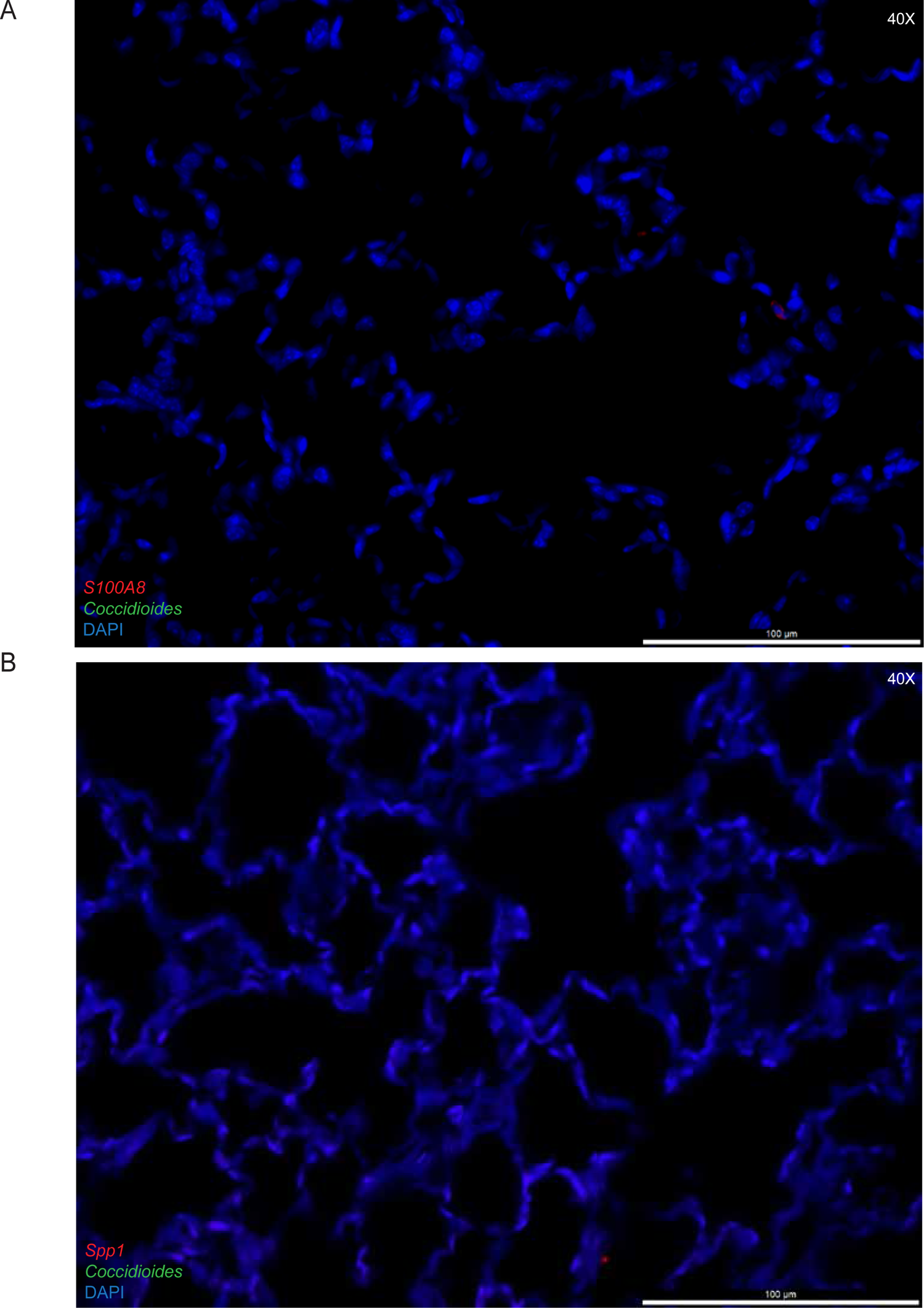
Uninfected IHCs. (A) Wild type mouse stained with S100A8 and DAPI, image **taken at 40X. (B) Wild type mouse stained with Spp1 and DAPI, image taken at 40X.**

**Supplemental Figure 5:**
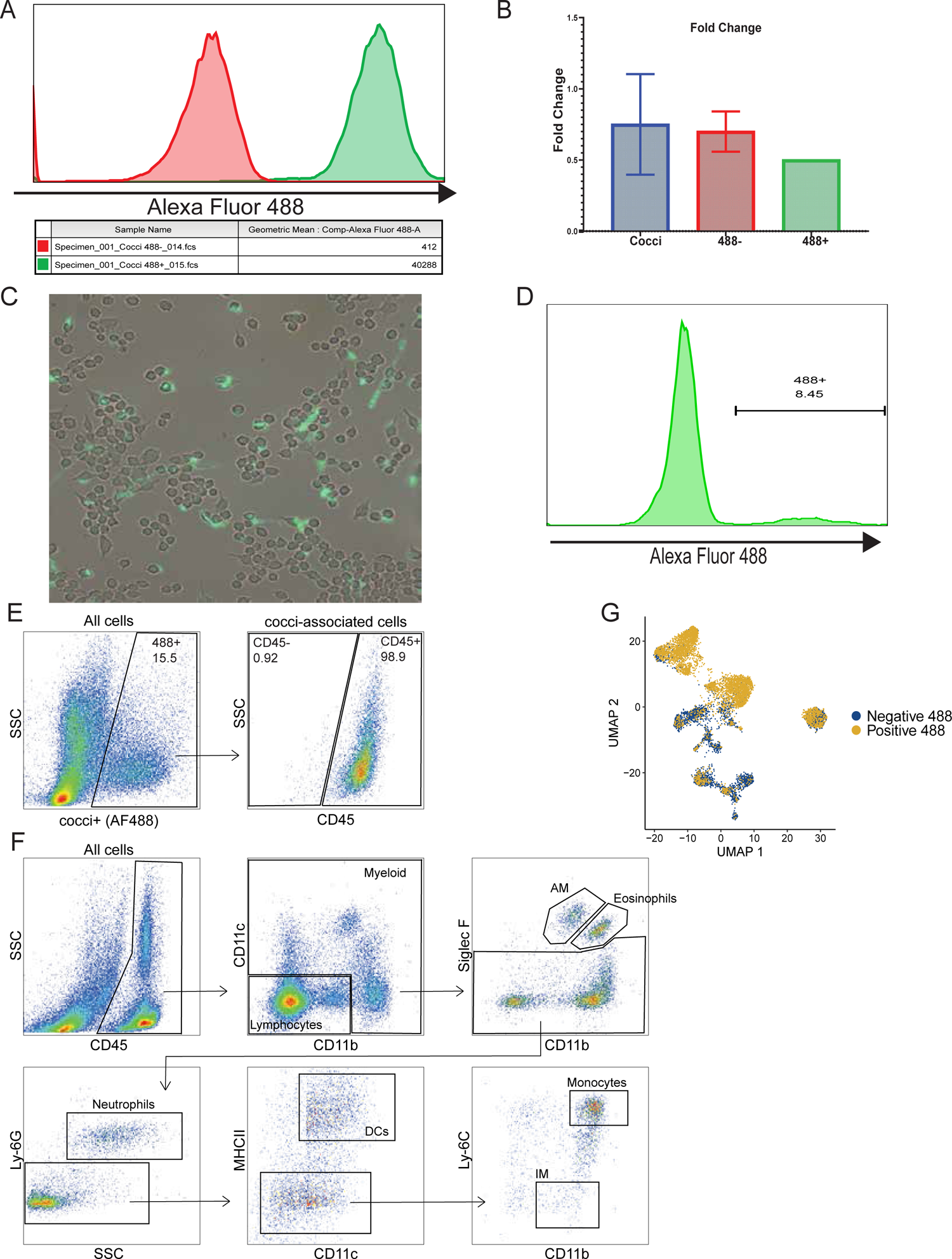
AF488 testing in vitro and flow gating strategy. Coccidioides was stained as described in the methods with AF488. (A) Flow cytometry analysis of labeled Coccidioides, AF488+ label. (B) Viability of labeled Coccidioides over time. 2X GYE were plated and CFU’s were counted after three days on triplicate plates. Error bars represent 2 experimental replicates. Statistical analysis was performed using a two-way ANOVA followed by Tukey’s multiple comparisons test. No statistical differences were found between the days per condition. (C) Labeled Coccidioides (green) co-cultured with 0.1 MOI for 2 hrs with RAW 264.7 macrophages (unlabeled) image taken at 20X. (D) Co-culture with AF488+ Coccidioides for 2 hrs at 0.1 MOI. Histogram: coculture gated on live AM. (E-F) Flow cytometry gating strategy for innate panel to identify immune cell populations. The first panel shows AF488+ from all cells and then is gated for CD45+ for infected samples. F shows the gating strategy for all cell types in an uninfected, AF488+/CD45+ gate that was applied after E gating strategy following the AF488 gating in the first panel.

